# Axonal spike-count regimes link spinal cord stimulation periodicity to artificial sensory detection and discrimination in rodents

**DOI:** 10.64898/2026.04.25.720749

**Authors:** Taylor J. Tvrdy, Aditi Bhattacharya, Rahul Radhakrishna, Jacob C. Slack, Amol P. Yadav

**Affiliations:** Lampe Joint Department of Biomedical Engineering, University of North Carolina at Chapel Hill and North Carolina State University, Chapel Hill, NC, USA; UNC School of Medicine, University of North Carolina at Chapel Hill, Chapel Hill, NC, USA; Max Planck Institute for Biological Intelligence, Seewiesen, Germany; UNC Neuroscience Center, University of North Carolina at Chapel Hill, Chapel Hill, NC, USA; UNC Neurosurgery, School of Medicine, Chapel Hill, NC, USA

**Keywords:** spinal cord stimulation, computational model, aperiodic stimulation, artificial sensations, axonal spikes

## Abstract

Electrical stimulation is widely used to evoke artificial sensation, yet how temporal stimulation patterns are transformed into neural activity and perception remains poorly understood. Recent behavioral studies have shown that increasing the aperiodicity of spinal cord stimulation pulse trains alters both detection thresholds and discrimination performance despite constant pulse counts, suggesting a role for temporal structure beyond rate alone. However, the neural mechanisms underlying these effects remain unresolved.

We developed a biophysically grounded computational framework linking a finite-element model of the rodent spinal cord, conductance-based axon simulations, and observer decision models to investigate how stimulation periodicity shapes neural responses and resulting behavior. By systematically varying stimulation amplitude, frequency, and inter-pulse interval variability, we identified distinct spike-count regimes arising from interactions between stimulation timing and axonal membrane dynamics. These regimes ranged from single spikes and small volleys to sustained spike trains, and they exhibited diverse frequency- and variability-dependent trends.

No single regime was able to sufficiently explain experimentally observed detection threshold trends; instead, mixtures of regimes accurately reproduced both frequency-dependent threshold behavior and trial-level variability. Extending this framework to periodicity discrimination, we show that features derived from regime mixtures contain sufficient information to recover behavioral psychometric curves. Furthermore, observer model results provide a mechanistic account of behavioral asymmetries depending on the periodicity of the reference stimulus: discrimination relative to periodic inputs relied on combined rate and timing evidence, whereas discrimination relative to aperiodic inputs was dominated by timing irregularity.

These results establish a mechanistic link between stimulation temporal structure, axonal spike generation, and perceptual behavior. This framework suggests that spinal cord stimulation does not operate within a single fixed neural regime but instead engages a spectrum of spike-count regimes whose mixtures shape perception. These findings have important implications for the design of biomimetic stimulation strategies, highlighting temporal patterning as a key dimension for controlling sensory outcomes.

## 1. Introduction

Electrical stimulation is widely used to artificially evoke sensorimotor signals through direct interfaces with the nervous system, such as intracortical microstimulation (ICMS) and spinal cord stimulation (SCS). Classical stimulation methods have largely been limited to modification of duration, amplitude, frequency, and bursting of stimulation pulses (Bjanes & Moritz, 2019; Grill, 2019). More recently, biomimetic ICMS patterns and waveforms have been successfully employed to restore sensation in brain-computer interface (BCI) applications (Bensmaia et al., 2023; Chandrasekaran et al., 2025; Flesher et al., 2016; Hughes et al., 2026; O’Doherty et al., 2012; Salas et al., 2018). Studies have shown that biomimetic ICMS improves perceptual quality by evoking neural activity that closely resembles natural brain dynamics (Hughes et al., 2026; Hughes & Kozai, 2023). A similar mechanistic understanding of how nontraditional SCS evokes sensory perception is not yet available. Moreover, whether modifying the temporal characteristics of SCS impacts the recruitment of neurons in supraspinal brain areas remains to be ascertained.

The temporal pattern of ascending neural activity plays a central role in encoding tactile information (Birznieks & Vickery, 2017; Mackevicius et al., 2012; Nakatani et al., 2021). Even when mean firing rate is held constant, variations in spike timing can alter perceived intensity and stimulus quality, indicating that temporal statistics carry perceptually relevant information beyond rate alone (Ng et al., 2018; Sharma et al., 2022). At higher centers of the somatosensory pathway, neurons in the dorsal column nuclei and cortex exhibit irregular firing patterns with non-zero coefficients of variation (CV) of the inter-spike interval, and both rate and timing features contribute to stimulus discrimination (Harvey et al., 2013; Shishido & Toda, 2017). These observations suggest that natural somatosensory processing relies on temporally structured yet variable neural activity rather than strictly periodic spike trains. Consequently, when designing biomimetic artificial stimulation patterns to evoke naturalistic sensation, the role of stimulation periodicity warrants consideration, as temporally patterned stimulation has been shown to influence both neural responses and perceptual outcomes (Birznieks & Vickery, 2017; Harvey et al., 2013; Mackevicius et al., 2012; O’Doherty et al., 2012; Slack et al., 2024).

To investigate whether the periodicity of epidural SCS pulse trains changed detection thresholds and just-noticeable differences (JND), Slack, et al. employed a two-alternative forced choice behavioral task in rats (Slack et al., 2024). They showed that detection thresholds decreased as CV of inter-pulse interval increased at lower frequencies (10, 20, 50 Hz), but not at higher frequencies (100, 200 Hz). Rats discriminated aperiodic stimulation trains from periodic stimulation trains at all tested frequencies with discrimination performance increasing as the difference in CV between the two stimulation trains increased. Furthermore, they noticed that maximally aperiodic pulse trains (CV = 1) had smaller JNDs than periodic pulse trains (CV = 0). Since the pulse trains across all CV values for a given frequency were charge-balanced (same number of pulses in 2 seconds), the behavioral differences in detection and discrimination were attributed to temporal characteristics independent of pulse count. However, a mechanistic understanding of how the temporal patterns impacted spiking behavior which subsequently led to perceptual differences in detection and discrimination is necessary.

As the stimulation pulse trains become more aperiodic, the distribution of inter-pulse intervals broadens, resulting in both longer and shorter intervals. If inter-pulse intervals are sufficiently short, temporal summation of graded potentials can contribute to action potential generation at lower input amplitudes, and the resulting axonal spike trains produced may be a subset of the input pulse trains. Additionally, shortening of inter-pulse intervals interacts with axonal membrane dynamics, leading to refractory period violations, afterpotential-mediated changes in excitability, and frequency-dependent reductions in spike generation (Bostock et al., 1998; Grill & Mortimer, 1994; Hughes et al., 2026; Joucla & Yvert, 2012; Kilgore & Bhadra, 2004; McIntyre et al., 2002; Rattay, 1999). Due to the relation of these phenomena to short inter-pulse intervals, we expect they are likely to impact the number of axonal spikes reproduced as input stimulation frequency and CV increase, producing different spike-count regime responses at the neural level despite a constant number of input pulses.

Further, the number of spikes underlying perceptual detection cannot be directly inferred from behaviorally determined thresholds alone. Prior studies across sensory modalities have shown that perception in response to brief visual (Hecht et al., 1942), auditory (Kiang et al., 1965), tactile (Vallbo & Johansson, 1976), and electrical stimulation (Yadav et al., 2021) stimuli can arise from relatively sparse neural activity, and that perceptual decisions can be influenced by small numbers of spikes (Parker & Newsome, 1998). These findings suggest that perception may be supported by a range of possible spike-count regimes, from single spikes or small volleys to more sustained firing patterns. Consequently, temporally structured electrical stimulation may produce a wide array of perceptual outcomes based on the potential spike-count regimes. It is therefore important to understand how neuronal membrane dynamics shape spike generation under these conditions to better control the percepts elicited by stimulation.

Given the potential for stimulation frequency and CV to induce different axonal spike-count regimes, along with the uncertainty in which regime drives perception, the present study aimed to explore these effects. We developed a detailed biophysical model of the rat spinal cord coupled to a McIntyre-Richardson-Grill (MRG) axon model (McIntyre et al., 2002) to investigate the neural consequences of epidural spinal stimulation periodicity *in silico*. By computationally recreating the experimental paradigm of Slack, et al. (Slack et al., 2024), we transformed stimulation pulse trains into axonal spike trains and quantified how input amplitude and temporal variability alter spike generation and reproducibility. We hypothesized that 1) interactions between stimulation characteristics and membrane dynamics produce distinct spike-count regimes that explain the frequency-dependent detection threshold trends, and 2) these spike-count regimes provide sufficient neural evidence to account for the difference in JNDs from periodic (CV = 0) and aperiodic (CV = 1) trains. Overall, we developed a mechanistic tool that allowed us to connect biophysical modelling of SCS to rodent perceptual behavior. We envision using this tool to test novel biomimetic SCS patterns, stimulation waveforms, and electrode configurations to understand how they influence sensory detection, perception, and discrimination behavior in animals and humans.

## 2. Methods

### 2.1 Experimental Overview

All animal procedures and experiments described in this study were approved by the University of North Carolina at Chapel Hill and Indiana University Institutional Animal Care and Use Committees. The study uses rat behavior data from Slack, et al. (Slack et al., 2024) where they investigated the effects of epidural SCS periodicity on detection threshold and periodicity discrimination in rodents by utilizing two-alternative forced choice (2AFC) tasks. CV values were incremented from 0 (periodic) to 1 (maximally aperiodic) in 0.1 steps, resulting in 11 unique CV values. Stimulation pulse trains for both detection threshold and discrimination were charged-balanced per-frequency such that all CV values for a given frequency contained the same number of stimulation pulses in the 2-second duration. Temporal patterns for pulse trains were derived from a random gamma-distribution sampling of inter-pulse intervals, which is discussed further in the ‘Stimulus Pattern Generation’ section below.

Rats were initially trained to perform 2AFC associating the left and right ports with ‘stim’ and ‘no stim’, respectively, before moving on to the detection threshold task where CV and amplitude were then randomized for ‘stim’ trials at a given frequency (10, 20, 50, 100, 200 Hz). For each frequency, 20 trials for all CV and amplitude combinations were completed, and detection thresholds were defined as the input amplitude corresponding to 75% correct performance from sigmoid fits. The results showed downward trends in detection threshold amplitudes as CV increased for 10, 20, and 50 Hz, but similar amplitudes across CV values for 100 and 200 Hz, and that amplitudes decreased for all CV values as frequency increased.

Rats then learned 2AFC associating the left and right ports with aperiodic (1 CV) and periodic (0 CV) pulse trains, respectively, before advancing to two discrimination task blocks. Trial blocks were separated by the chosen standard CV of 0 (std0) or 1 (std1), and performed for the given frequencies (20, 50, 100, 200 Hz). For std0 blocks, the right (periodic) port was held at a constant 0 CV while the left (aperiodic) port CV was randomized from 0.1 to 1 in increments of 0.1. For std1 blocks, the left (aperiodic) port was held at a constant 1 CV while the right (periodic) port CV was randomized from 0 to 0.9 in increments of 0.1. Stimulus amplitudes for both blocks and across all CV values were chosen as the individual rat’s 0 CV detection threshold and held constant. Discrimination performance was calculated for each CV value as the fraction of total correct trials per frequency and per block, and sigmoid psychometric curves were then fitted to the points. Results showed that rats were able to reliably discriminate between periodic and aperiodic stimuli across all tested frequencies, but the JND (point corresponding to 75% correct) in CV was smaller for std1 blocks than for std0 blocks.

### 2.2 Computational Pipeline Overview

To computationally recreate the experimental paradigm described above, we developed a biophysically accurate rodent spinal cord with which we applied electrical stimulation using electrodes of the same dimensions and spinal location as described in the experiment above. Following the simulation of spinal tissue electrical responses to a single bipolar, biphasic 1 µA stimulation pulse, we located a point of maximal cross-sectional activation in the dorsal column white matter and exported its longitudinal rostral-caudal electric potential across the length of the spinal cord (Figure 1a). These electric potential values were then scaled in amplitude and combined with 2-s temporal pulse patterns, which were similarly constructed by random samplings of a gamma-distribution of inter-pulse intervals (Figure 1b). The simulated 2-s electric potential trains were then linearly scaled and used as extracellular potentials to drive a single-axon MRG model, and the axonal responses were used to produce nine different spike-count regimes (Figure 1c), which are discussed in further detail in the ‘Spike-Count Regimes’ section below.

**Figure 1:**
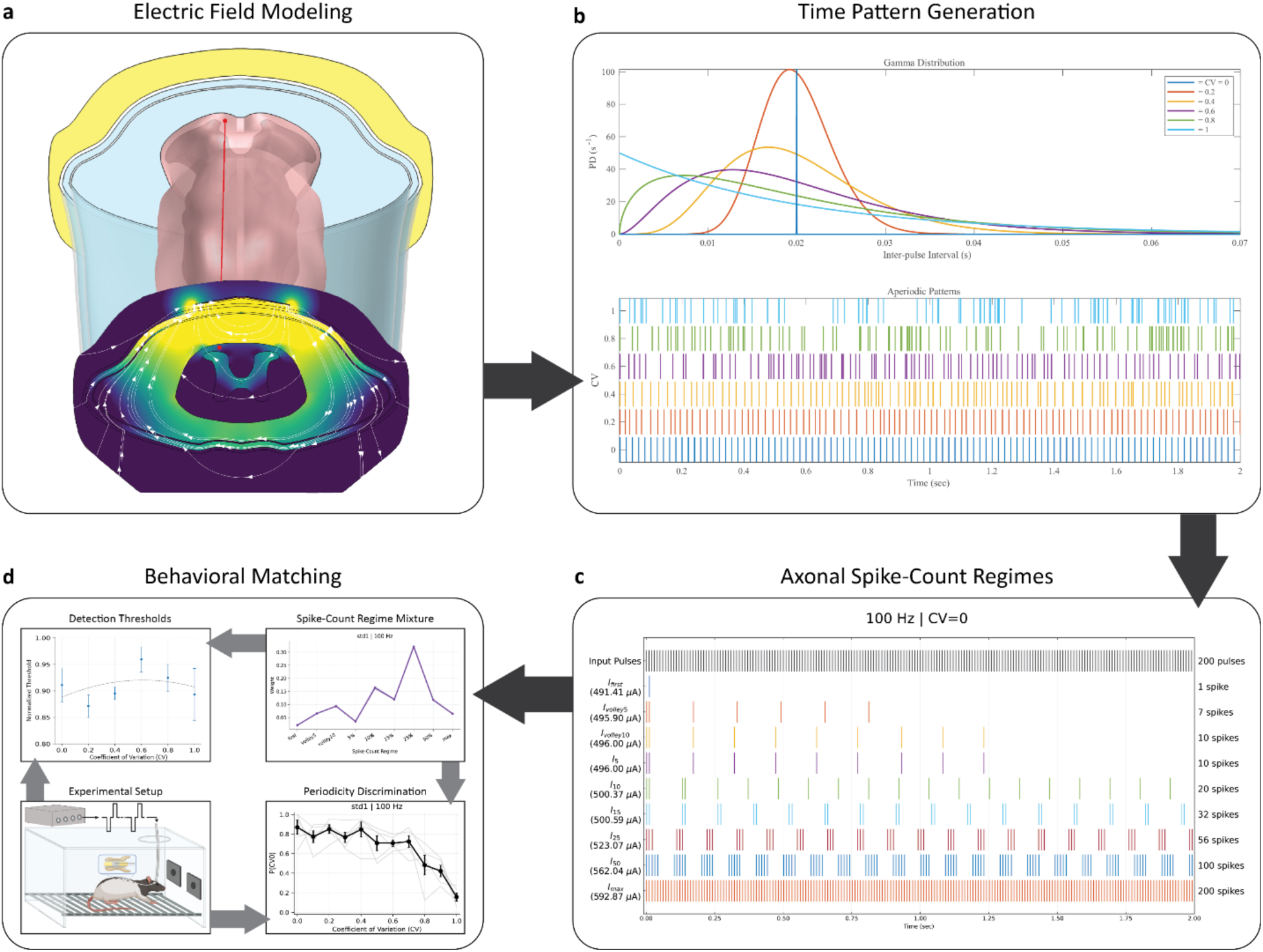
Computational Modeling Pipeline Overview. **a)** COMSOL biophysically accurate finite-element model of rodent spinal cord used to simulate spinal cord stimulation. The heatmap represents the cross-sectional current density at the rostral-caudal midline of the electrode in response to ±1 µA stimulation. A single axon trajectory (red) was placed in a dorsal column region of maximal current density, and electric potentials along its length were exported. **b)** Inter-pulse intervals were randomly sampled from a gamma distribution to produce pulse trains with the desired CV (0, 0.1, …, 1) and number of pulses (2*frequency) in a 2-s window. **c)** COMSOL-derived potentials in response to ±1 µA bipolar, biphasic stimulation were combined with generated time patterns and used as extracellular potentials to drive an MRG axon model. The ±1 µA input current amplitude was recursively scaled to satisfy the corresponding spike-count criteria, and this panel shows example spike-count threshold amplitudes and simulated axonal spike trains. **d)** Finally, mixtures of spike-count regimes were used to describe experimental observations of detection threshold and periodicity discrimination.

The various spike-count regimes were then used to investigate detection threshold and periodicity discrimination behavior (Figure 1d). Detection threshold analysis was aimed at exploring the capacity of individual spike-count regimes and learned mixtures of spike-count regimes to explain inconsistent trends across the tested frequency range. Periodicity discrimination analysis was conducted on the std0 and std1 blocks using observer models with learned mixtures of spike-count regimes to fit the behavioral psychometric curves based on the rate, timing, or hybrid (rate + timing) features of the regime mixtures.

### 2.3 Spinal Cord Model

We created a detailed model for rodent spinal cord SCS to provide a more precise understanding of the effects of injected currents on the underlying anatomical structures. To validate our model, the epidural electrode configuration used by Slack, et al. (Slack et al., 2024) was reproduced *in silico*, and the spinal cord activation profile was investigated. The Dana and Christopher Reeves Rodent Spine Atlas was used to construct the finite-element model (FEM) of the rat spinal cord in this study (Watson et al., 2009). Key anatomical features of epidural fat, dura mater, arachnoid mater, cerebrospinal fluid, pia mater, white matter, grey matter, and the central canal pictured on the atlas were scaled and traced using Apple Pencil 2 on an iPad Air 2020 in Shapr3D (Shapr3D, Zrt., n.d.). All the slices were then lofted, creating a comprehensive representation of each tissue. The various layers present in the model and their corresponding electrical conductivities acquired from previous modeling literature (Capogrosso et al., 2013, 2018; Khadka et al., 2020), can be seen in Supplementary Figure 1 and Supplementary Table 1. Stationary simulations were performed and therefore relative permittivity of the tissues was not taken into consideration. The spinal geometry from C1 to T5 was modeled to illustrate the response to epidural SCS with electrodes placed at the T4/T5 spinal level (Slack et al., 2024).

To simulate the experimental stimulation electrodes, two cuboids with the electrical properties of platinum (Pt) and dimensions of 1.5 x 1 x 0.25 mm were placed between the T4 and T5 spinal segments and mediolaterally centered on the midline of the spinal cord within the epidural fat layer. These electrodes were insulated with a custom material resembling properties of polydimethylsiloxane (PDMS). All sides of the electrodes were covered by the insulator except the stimulation surfaces that interfaced with the epidural fat layer. The electrical conductivities used for the electrodes and insulation can also be seen in Supplementary Table 1.

Computer-aided design (CAD) files of the spinal and electrode structures were individually imported into COMSOL 6.2 (COMSOL, Inc., n.d.) to form the composite model of non-overlapping domains for each structure. The pia mater was not given a thickness dimension but instead was represented by applying the meninges material properties to the outer surface of the white matter. An unstructured tetrahedral mesh of extra fine element size was used to mesh the entire model, resulting in 11,953,918 domain elements, 1,478,279 boundary elements, and 22,380 edge elements. The electrodes were simulated as boundary current sources and given opposite charge densities of ±1 µA to simulate bipolar, biphasic stimulation. A cylindrical infinite element domain surrounding the entire spinal cord structure was used as the electrical ground to enforce Dirichlet boundary conditions, and quasi-static state was assumed. Quadratic Lagrange elements were used for the scalar potential, and the resulting linear system was solved for 15,997,059 degrees of freedom using the Direct MUMPS solver (Capogrosso et al., 2018; Khadka et al., 2020).

After solving for the distribution of electric potential, a cross-sectional plane at the rostral-caudal midline of the stimulating electrodes was investigated to determine locations within the dorsal column white matter with the greatest cross-sectional current density, corresponding to areas likely to be excited at detection threshold (Figure 1a) (Holsheimer, 2002; Holsheimer & Buitenweg, 2014). Based on this investigation, the model was then embedded with a 3D cutline representing a single axon location and spanning approximately 25 mm from T5 to C1. The axon cutline was placed such that its trajectory was within the maximal area of activation while also residing in the white matter region along the length of the spinal cord to ensure no tissue-dependent changes in electric potential across its length.

The simulation was resolved after insertion of the axon cutline and electric potentials across the length of the axon were exported from COMSOL. As a result of the stationary solution used, the electric potential values in response to the ±1 µA stimulation were treated as instantaneous responses and were therefore linearly scalable. Although it is unlikely to experimentally activate only a single axon, we chose to model this extreme case to determine whether the behavioral trends could be explained by spike-count regimes with minimal neural recruitment, as *in vivo* experiments have shown similar capabilities from single neurons (Newsome et al., 1989; Parker & Newsome, 1998; Wang et al., 2007).

### 2.4 Stimulus Pattern Generation

The bipolar, biphasic aperiodic stimulation waveforms were then recreated similarly to the method of Slack et al. (Slack et al., 2024) (Figure 1b). Briefly, the pulse trains consisted of biphasic pulses with 200 µs phase duration and 50 µs inter-phase interval, for a total pulse duration of 450 µs. The inter-pulse intervals were then calculated for a 2-second pulse train using the gamma probability distribution function, *f*, described in equation (1) (O’Doherty et al., 2012)

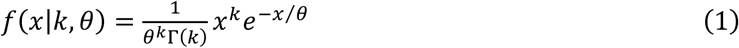

with inter-pulse interval *x* in milliseconds, gamma function Г, shape parameter *k*, and scale parameter θ. Minimum inter-pulse interval was enforced at the pulse duration of 450 µs to prevent overlapping pulses, and the degree of periodicity was modulated by changing the shape (*k*) and scale (θ) parameters based on the CV and frequency of the desired signal using equations (2) and (3).

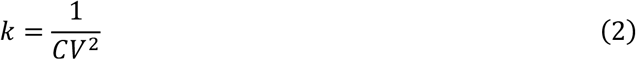

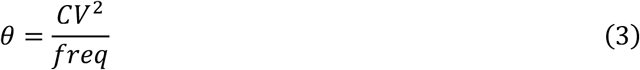

CV values from 0 to 1 in increments of 0.1 were tested; higher CV values produced a broader distribution of inter-pulse interval values, and thus higher aperiodicity. Frequency values of 10, 20, 50, 100, and 200 Hz were used and the 2-second pulse trains contained 20, 40, 100, 200, and 400 pulses, respectively, for each CV value. A 100-ms period of no stimulation was added to the beginning and end of each 2-s pulse train (2.2 seconds total) to ensure proper initialization and edge-spike handling in the MRG model. For reproducibility, the exact same pulse trains sampled from the gamma distribution during the experiments of Slack, et al. were used, as well as 99 new random samplings, resulting in 100 repetitions for each (CV, frequency) pair (11 CVs * 5 frequencies * 100 repetitions = 5,500 time patterns).

The axon cutline of electric potentials in response to a singular ±1 µA bipolar, biphasic pulse of SCS that were extracted from COMSOL were overlaid on the 5,500 generated time patterns to obtain space-time matrices of electric potentials, *V*_*e*_(*t*, *nodee*), to be used as 1 µA basis functions. Because the COMSOL solution was quasi-static, extracellular potentials were assumed to scale linearly with input current amplitude. Therefore, *V*_*e*_(*t*, *nodee*) basis functions were scaled to represent different input current amplitudes and used as extracellular potentials for the MRG axon model. The model was imported from ModelDB (McDougal et al., 2017) and the NEURON simulator (Hines & Carnevale, 1997) was interfaced within Python for further simulation. The axon was assumed to be cylindrical with 10 µm diameter, internodal length set to 100 times the diameter (Capogrosso et al., 2013), and all other parameters set to the default values of the original MRG model (Capogrosso et al., 2013; McIntyre et al., 2002), resulting in 26 simulated nodes of Ranvier. An interpolation function was used on the COMSOL-derived electric potential output to ensure accurate potential values at the x coordinates of the nodes, and the spiking behavior at the most rostral node was used for all subsequent analyses.

### 2.5 Spike-Count Regimes

As discussed previously, we expected that axonal membrane dynamics would produce varying spike-count responses to the stimulation trains due to short inter-pulse intervals as CV and frequency increase. Because it is unknown whether the rats detect the presence of a single spike, the full spike train, or an intermediate spike-count criterion (Holsheimer, 2002), we explored multiple spike-count regimes to determine whether specific operating ranges of the axon could describe the detection threshold and discrimination behavior observed by Slack, et al. (Slack et al., 2024).

For a given input current, *I*, let *N*_*spikkes*_(*I*) denote the number of axonal spikes produced in response to the 2-second pulse train, *N*_*pulses*_ = (2 · *freq*) denote the number of input stimulation pulses, *N*_*max*_ denote the maximum number of axonal spikes observed, and let *p* represent the five percentage-based spike-count criteria *p* ∈ {5%, 10%, 15%, 25%, 50%}. The nine spike-count regimes tested can thus be defined by equations (4a-e) describing the threshold amplitudes that satisfy the corresponding spike-count criteria.

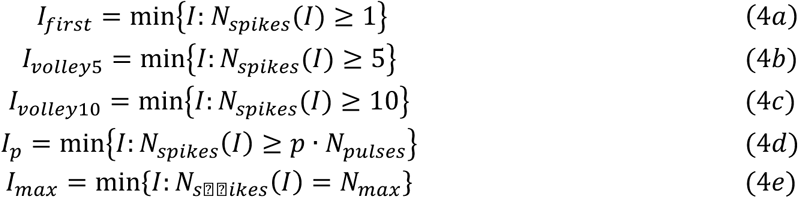

Although spike count generally increased with amplitude, refractory interactions at high CV and frequency values produced non-monotonic plateaus, and *N*_*max*_ did not always equal 100% of the input stimulation pulses. This was resolved by defining *I*_*max*_ as the minimum input current amplitude producing the maximum observed spike count, corresponding to the start of the plateau.

For each of the 5,500 time patterns, spike-count threshold amplitudes were computed by linearly scaling the 1 µA basis extracellular potentials, *V*_*e*_(*t*, *nodee*), and simulating the MRG axon model response. Amplitudes were bracketed up to a maximum search bound of 2 mA until the spike-count criterion was satisfied, followed by binary refinement to a tolerance of 0.1 µA. The number of spikes produced at the rostral terminal node were counted and used for subsequent analyses. Spike-count threshold amplitudes and the resulting number of axonal spikes across all frequencies and CVs can be seen in Supplementary Figures 2 and 3, respectively.

### 2.6 Detection Threshold Behavioral Matching

The identified spike-count regimes were then used to fit behavioral detection threshold data. Individual regimes were initially compared to the observed quadratic trends for 10, 20, 50, 100, and 200 Hz. Inaccurate fits from individual regimes led to the pursuit of spike-count regime mixtures that could better describe the experimental observations. Because the absolute amplitudes required to reach neighboring spike-count regimes were often tightly clustered, especially near CV = 0 (Supplementary Figure 2), small variability across rats, sessions, trials, or axon recruitment conditions could plausibly shift the active regime.

We therefore implemented spike-count regime mixtures for each CV value of a given frequency (per-point mixture) as a flexible approximation of the plausible regime variability, rather than as a claim that a fixed biological observer explicitly estimates a separate mixture at each CV/frequency point. Per-point mixtures weights were optimized at each CV, the resulting mixed threshold curve was normalized within frequency, and a bias term was fit to match the behavioral target. The per-point threshold mixtures were then compared against the single spike-count regimes to determine best-fits for the behavioral quadratic trends. Additionally, per-point mixtures were learned for the Slack-matched timing pattern repetition to fit the observed datapoints themselves, and the capacity to recover observation variability beyond the quadratic trends was assessed.

### 2.7 Periodicity Discrimination Behavioral Matching

To determine whether spike-count regimes provided sufficient neural evidence to account for behavioral periodicity discrimination, we developed a template-matching observer model linking simulation-derived features to the psychometric curves reported in Slack et al. (Slack et al., 2024). The observer model pipeline overview can be seen in Figure 2, and the sections below provide further detail about individual steps.

**Figure 2:**
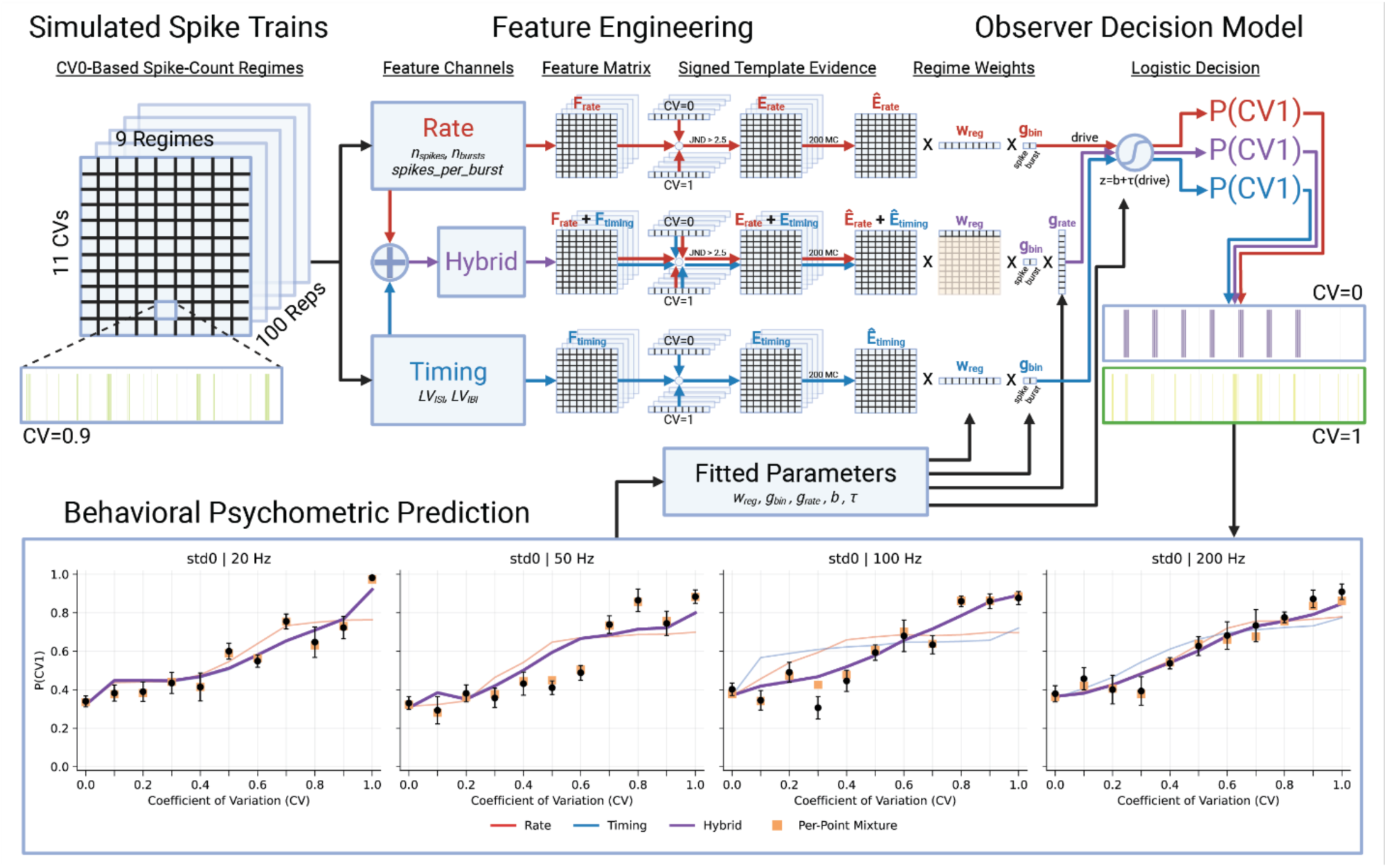
Periodicity Discrimination Model Overview. Simulated CV0-based spike trains were converted into rate, timing, and hybrid feature matrices (*F*_*rate*_, *F*_*timing*_, *F*_*rate*_ + *F*_*timing*_) for each frequency. For each model family, signed evidence relative to the CV = 0 and CV = 1 template endpoints (*E*_*rate*_, *E*_*timing*_, *E*_*rate*_ + *E*_*timing*_) was computed and normalized separately for each Monte Carlo training set (*Ê*_*rate*_, *Ê*_*timing*_, *Ê*_*rate*_ + *Ê*_*timing*_), then combined using per-block (same mixture for each CV and frequency in the same block) or per-point (hybrid only; separate mixtures per-CV, per-frequency) regime mixture weights (*w_reg_*). The 10-ms and 1-ms evidence channels were retained separately and integrated with fitted gains (*g*_*bin*_) before logistic readout; in the hybrid model, the rate pathway included an additional fitted rate-gain (*g*_*rate*_) term to allow different relative timing and rate contribution across CV values. The resulting drive signals were passed through a logistic decision stage (*z* = *b* + τ(*drive*)) to predict the psychometric curves. Parameters (*w*_*re*g_, *g*_*bin*_, *g*_*rate*_, *b*, τ) were optimized to minimize mismatch between predicted and observed psychometric curves. The lower panels show example behavioral datapoint (black) predictions for the per-block rate (red), timing (blue), and hybrid (purple) models, as well as the per-point hybrid model (orange).

#### 2.7.1 Simulated Spike Trains

In the experimental paradigm, discrimination stimuli were delivered at each rat’s 0 CV detection threshold for a given frequency and held constant across other CV values. To reproduce this structure computationally, we first computed the nine spike-count threshold amplitudes at CV = 0 for each (frequency, repetition) pair and then simulated the axonal responses to those constant input amplitudes for the corresponding frequency- and repetition-matched time patterns at all other CV values. This ensured that discrimination evidence was evaluated under fixed-amplitude conditions consistent with the behavioral task. Resulting spike counts for all frequencies and CVs in response to the CV0-based threshold amplitudes can be seen in Supplementary Figure 4.

#### 2.7.2 Feature Engineering

For each simulated spike train response indexed by frequency, CV, spike-count regime, and repetition, spike times were analyzed at two temporal resolutions to capture both fine and coarse temporal structure (Mackevicius et al., 2012; Samonds & Bonds, 2004; Victor & Purpura, 1997; Wang et al., 2007). Spike times were binned at 1-ms resolution to preserve individual spikes (Supplementary Figure 5a) and at 10-ms resolution to capture burst-level structure (Supplementary Figure 5b), consistent with previously observed temporal acuity findings from *in vivo* neural recordings (Mackevicius et al., 2012; Nemenman et al., 2008; Samonds & Bonds, 2004; Victor & Purpura, 1997; Wang et al., 2007). Contiguous active 10-ms bins were collapsed into single burst events defined by the first and last spike times within each event. From these spike- and burst-based representations, rate, timing, and hybrid (rate + timing) feature channels were constructed and later converted into signed template evidence relative to the CV = 0 and CV = 1 endpoint templates.

Rate features were calculated over each 2-s window as the number of spikes (*n*_*spikkes*_), the number of burst events (*n*_*bursts*_), and the mean number of spikes per burst (*spikkes*_*per*_*burst*), thereby capturing overall spike count and burst intensity (Supplementary Figure 5c). Timing features were quantified using the local variation (LV) of inter-spike intervals (*LV*_*IBI*_) for the 1-ms representation and the LV of silence intervals between burst events (*LV*_*IBI*_) for the 10-ms representation, thereby measuring temporal irregularity of spiking and bursting activity (Supplementary Figure 5d) (Birznieks C Vickery, 2017; Ng et al., 2018; Shinomoto et al., 2005). Other spike-train distance measures were also evaluated (Cutts C Eglen, 2014; Satuvuori & Kreuz, 2018; van Rossum, 2001; Victor & Purpura, 1997; Wang et al., 2007), but they showed regime-shift sensitivity from periodic (CV = 0) to weakly aperiodic (CV = 0.1) trains that was not present in the behavioral data; LV was therefore selected as a timing metric that is relatively insensitive to overall rate, described by equation (5) for consecutive silence intervals (*T*) (Shinomoto et al., 2005).

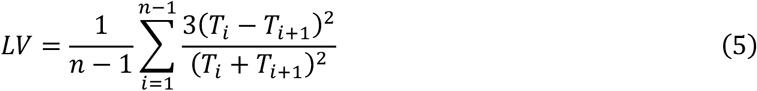

Because rats were initially trained to discriminate between strictly periodic (0 CV) and maximally aperiodic (1 CV) stimulation, repetition-matched templates were constructed at CV = 0 and CV = 1 for each frequency and spike-count regime pair. These endpoint templates served as internal reference anchors against which intermediate-CV stimuli were evaluated. Rate evidence was computed using a Weber-like JND criterion (Ekman, 1959; Weber, 1834). Specifically, for a candidate rate feature value, *f*_*r*_, and template rate feature value, *t*_*r*_, the two were considered indistinguishable if max (*f*_*r*_, *t*_*r*_)⁄min (*f*_*r*_, *t*_*r*_) ≤ *k_JND_*, where *k*_*JND*_ = 2.5 was chosen conservatively based on previously reported frequency-discrimination limits in animal models, corresponding to a minimum observed *k*_*JND*_ of approximately 2.7 (Yadav et al., 2021). Each rate feature (*n*_*spikkes*_, *n*_*bursts*_, spik*kes*_*per*_*burst*) contributed a vote of -1 if only the 0 CV template was within JND, +1 if only the 1 CV template was within JND, and 0 otherwise; rate evidence was defined as the mean of non-zero votes (Supplementary Figure 5e). Timing evidence was computed as the signed difference in absolute deviation of the timing features (*LV*_*IBI*_, *LV*_*IBI*_) from the two endpoint templates, such that positive values indicated closer similarity to the CV = 1 template (Supplementary Figure 5f). These repetition-level rate and timing evidence values were then endpoint-normalized and passed to the observer decision model to fit the behavioral psychometric curves.

#### 2.7.3 Observer Decision Model

The observer decision model was evaluated using 200 Monte Carlo 80/20 train/test splits over the simulated repetitions for each frequency and spike-count regime condition. For each split, 80% of repetitions were assigned to the training set and used to construct mean endpoint-normalized rate and timing evidence curves across CVs for both the spike (1-ms) and burst (10-ms) representations (Supplementary Figure 6), while the remaining 20% were held out for testing. The same endpoint normalization defined by the training repetitions was then applied to the held-out repetitions to generate test-set evidence curves, allowing generalization to unseen simulated responses to be assessed. For rate curves, an additional reliability factor was applied at each CV, defined as the fraction of repetitions producing a non-zero JND vote. This step down-weighted rate signals that were inconsistent across repetitions while preserving reproducible rate differences.

Within each model family (rate, timing, hybrid), normalized evidence was combined using a learned convex mixture over the nine CV0-based spike-count regimes. Non-negative per-block regime weights summing to one were optimized separately for the std0 and std1 blocks, such that a single weight vector was shared across all frequencies and CVs within a block to emphasize mechanistic differences between the two blocks. After this regime-mixture stage, the 1-ms spike evidence and 10-ms burst evidence were retained as separate channels and scaled by fitted gains before being summed into a latent decision drive, allowing the observer to learn how to relatively weigh the different evidence streams. The hybrid model contained an additional stage where an optimized rate-gain parameter was applied to the rate evidence streams, allowing the model to adjust the relative rate and timing contribution across CV values. Additionally, a per-point hybrid model was constructed similarly to the per-block hybrid model, except with separate mixtures for each CV and frequency point (per-point mixtures). The per-point hybrid model was therefore treated as a less-constrained comparison model that asked whether plausible shifts among spike-count regimes could recover local behavioral datapoint structure when mixture weights were allowed to vary by CV and frequency.

The pooled latent drive signals for each model were then mapped to the probability of choosing the aperiodic stimulus, *p*(*CV*1), through a logistic decision rule, *z* = *b* + τ(*drive*). For each of the per-block and per-point models, downstream decision parameters were optimized separately for each frequency-block condition, including the logistic intercept (*b*), logistic sensitivity term (τ), 1-ms and 10-ms channel gains, and the hybrid rate-gain. The resulting logistic output provided the predicted psychometric curve for that panel. Parameters were fit by minimizing training set error relative to the animal behavioral psychometric datapoints from Slack, et al. (Slack et al., 2024), and were then evaluated on the held-out test-set evidence curves.

## 3. Results

Using our computational modeling pipeline integrating simulated SCS electric potentials, MRG model axon responses, and behavioral observation targets, we were able to reproduce a SCS periodicity protocol *in silico*. Various axonal operating ranges were reliably reproduced with changes in input current amplitude, and mixtures of these spike-count regimes were able to describe both detection threshold and periodicity discrimination behavior. This work provides mechanistic insight into axon spiking dynamics in response to SCS and how they can shape behavior.

### 3.1 Membrane Dynamics Produce Distinct Spike-Count Regimes

Small changes in input amplitude produced distinct spike-count regimes in the simulated axon. Based on the spike-count criteria tested, we show that single spikes, small spike volleys (5 – 10 spikes), partial sustained trains, and near-complete spike trains are all possible phenomena reaching the brain in response to 2-s trains of spinal cord stimulation. The detection threshold amplitudes observed by Slack, et al. (Slack et al., 2024) had a peak value of about 370 µA, and the mean amplitudes for our simulated spike-count thresholds range from approximately 450 µA for *I*_*first*_ to 725 µA for *I*_*max*_ (Supplementary Figure 2). While these values don’t precisely overlap, the small discrepancies are acceptable given the model simplifications and axon placement at a non-maximal location in the simulated electric field (Figure 1a) to ensure the axon trajectory remained in the white matter along the entire length of the spinal cord.

The investigated spike-count regimes present different trends across frequencies and CVs due to the varying effects of short inter-pulse intervals (Supplementary Figures 2 and 3). In general, as the spike-count criterion increases, the relation between threshold amplitude and CV initially trends downward (e.g., *I*_*first*_, Supplementary Figure 2), flattens out, and eventually trends upward (e.g., *I*_*max*_, Supplementary Figure 2), but the trend strength and crossover point differ across frequency. Downward trends for lower spike-count regimes arise from the temporal summation of short inter-pulse intervals at higher CVs, which requires less input current to produce spikes. Conversely, upward trends for higher spike-count regimes (*I*_*max*_ in particular) require greater amplitudes to overcome relative refractory violations, and absolute refractory violations (less than ∼1.4 ms for current MRG model settings) lead to *N*_*max*_ < *N*_*pulses*_ as frequency and CV increase (Supplementary Figure 7).

### 3.2 Detection Threshold Trends Emerge from Spike-Count Regime Mixtures

Absolute (Figure 3a) and normalized (Figure 3b) spike-count threshold amplitudes for the 10, 20, 50, 100, and 200 Hz frequencies were used to compare the capacity of single spike-count regimes and per-point mixtures to fit the corresponding experimental detection threshold behavioral data (Figure 3c). The single-regime comparisons used normalized thresholds from Figure 3b with a learned bias term to account for the vertical offset due to experimental data being normalized across rats. Per-point mixtures for each frequency used learned mixtures of absolute thresholds from Figure 3a by iteratively weighting the regimes at each CV independently, normalizing all CVs together, and updating the mixture weights until the error between the predicted and behavioral quadratic trends was minimized. Similarly, this process was repeated for the Slack-matched time pattern repetition and individual detection threshold datapoints were fit instead of quadratic trends.

**Figure 3:**
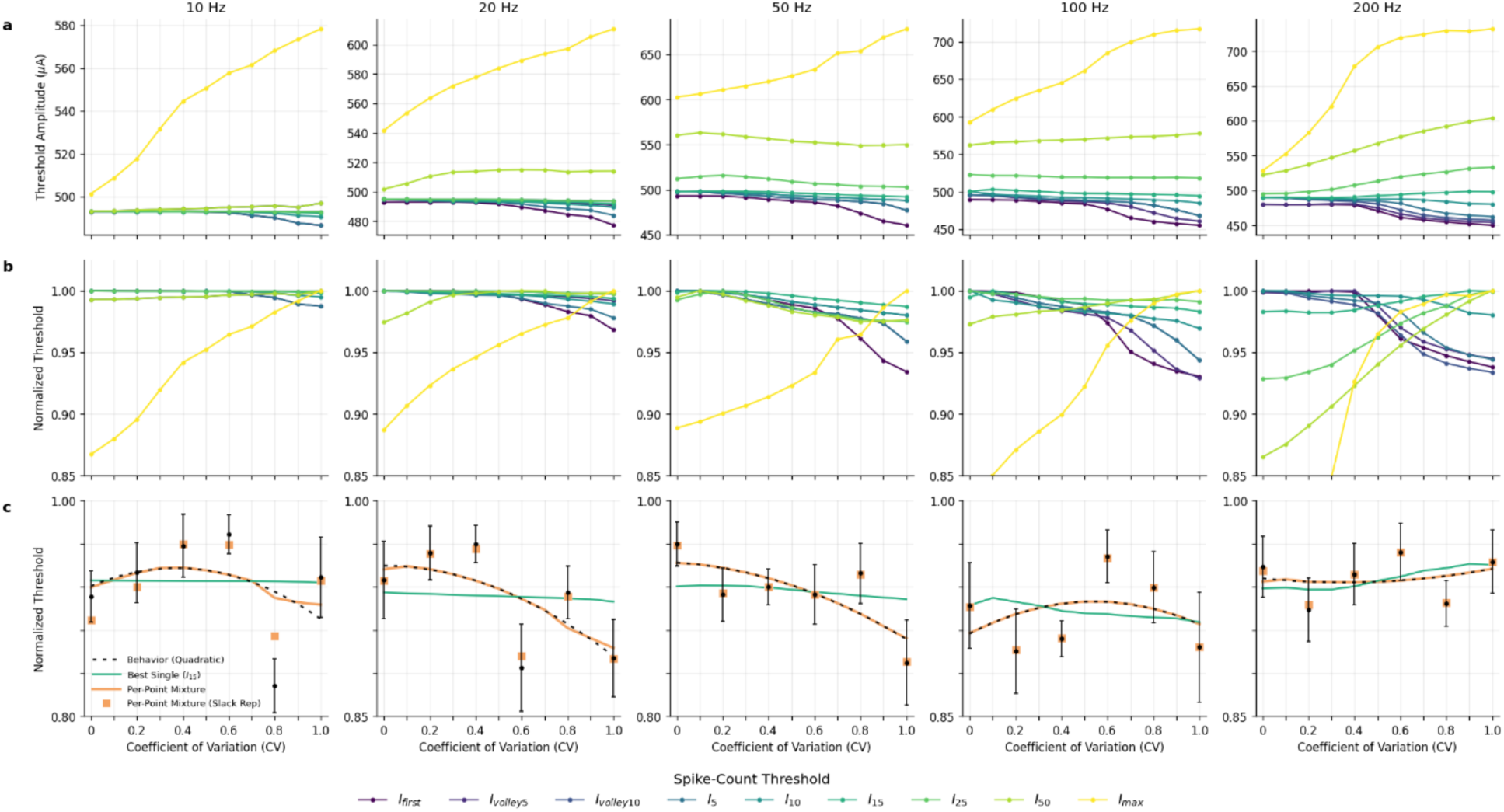
Detection Threshold Behavioral Matching. This figure shows simulated spike-count threshold amplitudes and compares them to the detection threshold trends observed experimentally. Colored traces represent the different spike-count thresholds marked in the bottom legend. **a)** Absolute threshold type amplitudes (µA) as a function of CV. **b)** Normalized threshold type amplitudes as a function of CV. Values normalized by the maximum value across all CVs for each threshold type individually to illustrate their directional trends. **c)** Normalized behavioral threshold datapoints (mean ± SEM) were fit with quadratic trends, and simulated spike-count regimes were used to match the trends. The best-fit single regime (*I*_15_; 0.012 mean RMSE) could not accurately describe the observed behavior, but per-point regime mixtures could describe behavioral trends (all simulated repetitions; 0.001 mean RMSE) and individual datapoints (Slack-matched time pattern repetition; 0.006 mean RMSE).

As CV increases, the behavioral quadratic fits in Figure 3c show inconsistent trends with downward parabolic curves for 10 and 100 Hz, an upward parabolic curve for 200 Hz, and monotonic decreasing trends for 20 and 50 Hz. Even the best-fit single spike-count regime was unable to accurately capture these trends across frequencies (*I*_15_; 0.012 mean RMSE), but the per-point mixtures using all repetitions did (0.001 mean RMSE) capture it. Further, when using the Slack-matched time pattern alone, it was possible to accurately reproduce the individual datapoint variability beyond just the trends using per-point mixtures (0.006 mean RMSE). These results illustrate that single spike-count regimes cannot describe detection threshold trends, and that a mixture of regimes is more likely.

While the per-point mixtures have additional degrees of freedom they should be interpreted as flexible upper-bound fits rather than uniquely identifiable biological mechanisms. As mentioned previously, the tight clustering in spike-count threshold amplitudes (Figure 3a) could conceivably lead to variability in axon operating ranges across rats, days, and trials. Additionally, the weights attributed to the two independent per-point mixture fits for the behavioral quadratic trends (all repetitions) and individual datapoints (Slack repetition) were qualitatively similar (Supplementary Figure 8). For example, at low CV values for 10 and 20 Hz, both mixtures identified strong *I*_*max*_contribution to account for the upward behavioral trends at these points (Figure 3c).

### 3.3 Observer Models Reproduce Behavioral Discrimination

To determine whether spike-count regimes provide sufficient neural evidence to account for behavioral periodicity discrimination, we evaluated observer models using rate, timing, and hybrid (rate + timing) channels derived from the simulated spike trains. For each model, evidence from the nine CV0-based spike-count regimes was combined using learned convex mixture weights and passed through a logistic decision node to generate predicted psychometric curves. Per-block rate, timing, and hybrid models were fit with spike-count regime mixtures across frequencies within each trial block (std0 and std1) to provide insight into mechanistic block differences, while the per-point hybrid model utilized independent mixtures for each datapoint to offer an experimentally feasible upper-bound in performance. All models were evaluated using 200 Monte Carlo train/test splits over the simulated repetitions and performance was compared using held-out test-sets.

All three per-block models were able to reproduce the monotonic structure of the behavioral psychometric curves, and the per-point model was able to recover individual datapoint variability (Figure 4). Block-averaged RMSE values for the per-point hybrid model revealed upper-bound fits (RMSE = 0.031) for both blocks, and paired Wilcoxon tests across Monte Carlo draws of the per-block models showed std0 and std1 blocks were best fit by the hybrid (RMSE = 0.074) and timing (RMSE = 0.094) models, respectively (Supplementary Table 2) [50]. These results indicate that spike-train features derived from the mixtures of different simulated axonal response regimes contain sufficient information to reproduce the behavioral discrimination patterns. Learned spike-count regime mixtures for the per-block and per-point models can be seen in Supplementary Figure 9.

**Figure 4:**
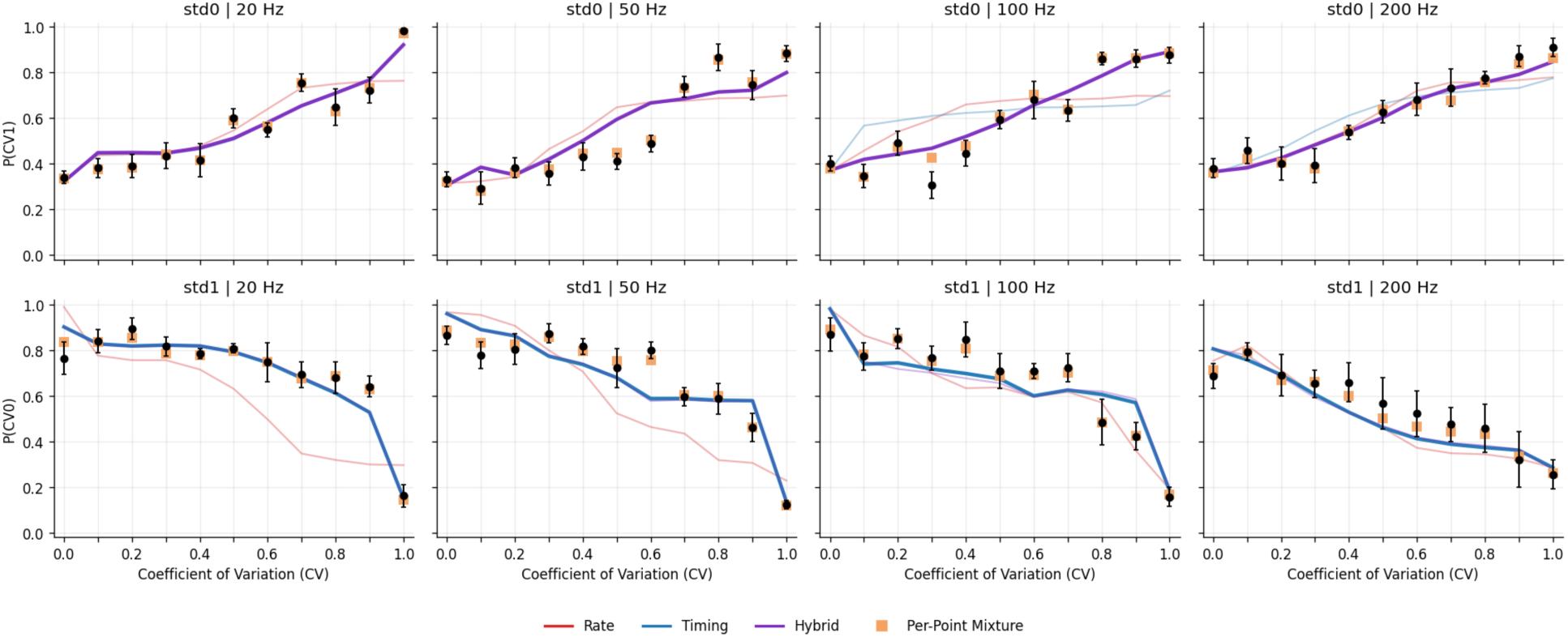
Periodicity Discrimination Behavioral Matching. This figure shows the resulting model fits to the experimentally observed psychometric curves. Individual behavioral datapoints (black) show the mean ± SEM and the colors represent the per-block mixture rate (red), timing (blue), and hybrid (purple) models, and the per-point mixture hybrid model (orange), as described in the bottom legend. Per-block models were fit across all CVs and frequencies in each block (std0 and std1) with a shared mixture of CV0-based spike-count regimes, while the per-point model had separate mixtures for each CV and frequency. The best-fit per-block models are shown in bold.

### 3.4 Discrimination Mechanisms Differ Between std0 and std1 Blocks

Inspection of the per-block model fits revealed distinct discrimination mechanisms for the two experimental trial blocks. In the std0 block, where the periodic stimulus (CV = 0) served as the reference, the hybrid model significantly outperformed both the rate (45.68% worse mean RMSE, p < 0.001) and timing (42.36% worse mean RMSE, p < 0.001) models (Supplementary Table 2), determined by paired Wilcoxon tests across Monte Carlo draws. Investigation of the learned spike-count regime mixtures revealed relatively equal weights for all regimes (Supplementary Figure 9a). This result indicates that discrimination relative to a fully periodic reference train relies on a combination of rate and timing information.

Conversely, in the std1 block, where the maximally aperiodic stimulus (CV = 1) served as the reference, the per-block timing model significantly outperformed both the rate (63.82% worse mean RMSE, p < 0.001) and hybrid (3.03% worse mean RMSE, p < 0.001) models, determined by paired Wilcoxon tests across Monte Carlo draws. Although the timing model consistently outperformed the hybrid model across Monte Carlo draws, their performance was similar (Supplementary Table 2), and their learned spike-count regime mixtures were nearly identical (Supplementary Figure 9a). This result indicates that discrimination relative to maximally aperiodic reference trains relies primarily on spike and burst timing irregularity as opposed to overall spike count, burst count, or burst intensity.

To further support this claim, the effective rate shares for the per-block and per-point hybrid models were calculated as the proportion of total model drive attributable to rate evidence after applying the learned rate/timing scaling (Figure 5a). The mean effective rate share was then calculated using the areas under the curves (AUC) to provide a summary metric of average rate-reliance across CV (Figure 5b). Per-block and per-point hybrid models displayed significant increases in rate-reliance as frequency increased for both std0 and std1 blocks (Figure 5b; all significance markers represent Bonferroni-corrected p < 0.05). The per-point hybrid model showed rate-reliance in both blocks and did not exhibit a significant pooled std0/std1 block difference (p = 0.069), whereas the per-block hybrid model showed significantly greater rate-reliance in std0 than std1 (p < 0.001). The low rate-reliance of the std1 per-block hybrid model paired with its regime mixture similarity to the std1 per- block timing model render the two models practically identical, and explains the 3.03% difference in RMSE performance. The differences in rate-reliance between the best-fit std0 and std1 per-block models support our second hypothesis, and we propose that the experimental JND asymmetry reflects block-dependent differences in the reliance of rate and timing evidence.

**Figure 5:**
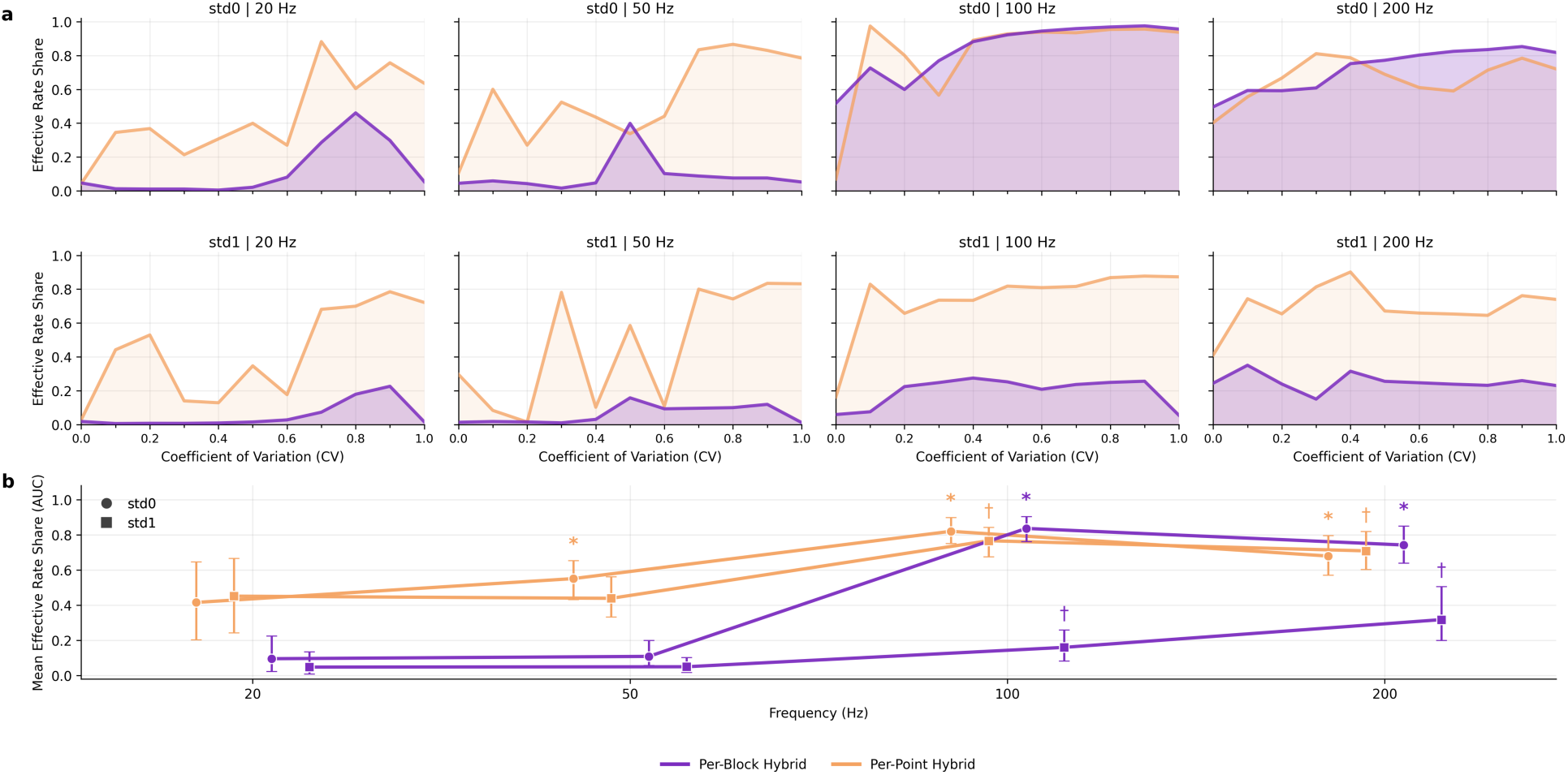
Rate Reliance of Hybrid Discrimination Models. This figure shows the effective rate share for the per-block and per-point hybrid models as described by the bottom legend. **a)** Bold traces show effective rate share, calculated as the proportion of total model drive attributable to rate evidence after learned rate/timing scaling. Shaded areas under the curves illustrate the AUC used as a summary metric of average rate reliance across CV. **b)** Mean effective rate share AUC values are shown for each frequency. Circle datapoints and asterisk significance markers denote std0 blocks, whereas square datapoints and dagger significance markers denote std1 blocks. Significance markers indicate Bonferroni-corrected p < 0.05 for comparisons against the corresponding 20 Hz AUC value. Error bars show [2.5, 97.5] percentile intervals, indicating cross-repetition spread.

## 4. Discussion

The biophysically grounded spinal cord model developed in this study provides a mechanistic bridge between stimulation parameters, axonal dynamics, and behavioral outcomes. By linking finite-element electric field simulations with MRG axon modeling and observer decision models, we demonstrate that experimentally obtained detection threshold trends and periodicity discrimination behavior can be explained by non-linear axonal responses to input stimuli. This establishes a framework for interpreting behavioral responses to SCS in terms of stimulation-driven axonal operating regimes and their downstream rate- and timing-based decision evidence and offers a pathway toward more principled design of stimulation paradigms.

A central finding of this work is that interactions between stimulation timing and axonal membrane dynamics give rise to distinct spike-count regimes that vary systematically with frequency and CV. These regimes emerge from well-known nonlinearities in axonal excitability, including temporal summation and refractory effects, which together shape how input pulse trains are transformed into spike outputs (Bostock et al., 1998; Grill & Mortimer, 1994; Joucla & Yvert, 2012; Kilgore & Bhadra, 2004; McIntyre et al., 2002; Rattay, 1999). Lower spike-count regimes benefit from short inter-pulse intervals through temporal summation, while higher spike-count regimes are increasingly constrained by refractory violations, limiting spike reproducibility at higher frequencies and CVs. Although the regimes explored here were derived from a single axon, axons of different diameters and spatial locations would likely occupy multiple operating modes simultaneously *in vivo*. Thus, the behavioral mixture-based fits observed here may instead reflect aggregate recruitment across multiple axonal operating regimes (Bensmaia et al., 2023; Mackevicius et al., 2012).

No single regime was sufficient to account for the full range of observed detection threshold trends. Instead, per-point mixtures of regimes were able to reproduce both the frequency-specific behavioral trends and the Slack-matched datapoint variability. These mixtures should not be interpreted as literal biological estimates of CV-specific regime weights, but rather as flexible upper-bound approximations of plausible variability in the active axonal operating regime. This interpretation is supported by the tight clustering of absolute threshold amplitudes across neighboring regimes (Figure 3a), especially near CV = 0, where small changes across rats, days, and trials could plausibly shift which regimes are expressed. This interpretation highlights an important consideration for stimulation studies: detection thresholds may not correspond to a single fixed spike-count regime but rather reflect probabilistic engagement of multiple regimes depending on both the stimulation conditions and underlying membrane dynamics.

Extending beyond detection thresholds, our observer modeling results demonstrate that these spike-count regimes contain sufficient neural information to reproduce behavioral periodicity discrimination. The per-point hybrid model showed that allowing regime mixtures to vary locally by CV and frequency could recover finer behavioral datapoint structure, whereas the per-block models provided a more constrained test of shared underlying mechanisms. The per-block model results demonstrated that the relative contribution of rate and timing features depends on the structure of the reference stimulus. In the std0 condition, where periodic stimulation served as reference, discrimination was best explained by the per-block hybrid model incorporating both rate and timing features. In contrast, in the std1 condition, where aperiodic stimulation served as the reference, discrimination was more accurately captured by the per-block timing model. This shift in feature reliance is consistent with a mechanistic interpretation for the asymmetric JNDs observed experimentally, with greater sensitivity to deviations from aperiodic stimuli arising from increased sensitivity to spike-timing irregularity relative to rate differences (Arabzadeh et al., 2006; Mackevicius et al., 2012; Wang et al., 2007).

To illustrate these mechanisms near the two CV endpoints, consider the periodic reference stimulus (CV = 0), which provides a highly regular timing template. Small increases in temporal irregularity for nearby CVs may produce limited perceptual separation, whereas larger CV differences increasingly alter spike generation and reproducibility through short-interval summation and refractory effects, creating rate-relevant evidence. In contrast, for the aperiodic reference stimulus (CV = 1), nearby CVs still remain highly irregular, even as CV decreases modestly, making discrimination more naturally supported by timing-based differences. The block-specific feature reliance produced experimentally may therefore arise from the different rate-versus timing-based entrainment reinforced by the std0 and std1 blocks, respectively.

From an engineering perspective, these results have direct implications for the design of stimulation protocols. Modulating temporal structure to exploit membrane dynamics may allow for selective control over spike generation and reproducibility. For example, short inter-pulse intervals can be used to lower activation thresholds through temporal summation, potentially reducing required stimulation amplitudes. Conversely, avoiding refractory violations may be necessary when the goal is to preserve faithful transmission of intended spike patterns. More generally, the finding that spike-count regimes across CVs emphasize different neural features suggests that stimulation strategies could be designed to bias downstream evidence toward rate-based or timing-based representations, depending on the desired perceptual outcome (Bensmaia et al., 2023; Ng et al., 2018; Samonds & Bonds, 2004). Further, with well-documented frequency limitations due to Weber’s law (Bjanes & Moritz, 2019; Ekman, 1959; Yadav et al., 2021), utilizing timing-based encoding may provide increased bandwidth to the signals transmitted along axons (Arabzadeh et al., 2006; Nemenman et al., 2008).

Despite these insights, several limitations about the modeling conditions should be considered. The current model does not include dorsal or ventral roots, axon populations, or synaptic processing, which could influence activation mapping, active spike-count regimes, and downstream interpretation. Behavioral discrimination was also modeled using an idealized observer framework, which does not explicitly account for higher-order decision processes. Additionally, the per-point mixture models are intentionally flexible and should be interpreted as upper-bound demonstrations of information sufficiency rather than uniquely identified biological mechanisms – multiple combinations of axonal regimes could likely produce similar behavioral fits, and future work with population-level recordings or more anatomically detailed recruitment models would be needed to constrain these mixtures directly. In addition, statistical comparisons based on simulated repetitions and Monte Carlo train/test splits quantify robustness to time-pattern sampling and model fitting variability, but they should not be interpreted as animal-level population inference.

## 5. Conclusion

In conclusion, this study demonstrates that interactions between spinal cord stimulation timing and axonal membrane dynamics can produce distinct spike-count regimes that help explain both detection threshold trends and periodicity discrimination behavior. Per-point regime mixtures captured frequency- and CV-specific behavioral variability, while more constrained per-block observer models revealed that discrimination depends on block-specific weighting of rate and timing evidence. Together, these findings provide a mechanistic link between stimulation patterns, axonal spike generation, and behavioral outcomes. This modeling framework therefore acts as a tool that can be used to test how biomimetic stimulation patterns, waveforms, electrode configurations, and recruited axon populations may shape sensory detection and discrimination before experimental implementation.

## Figures

**Supplementary Figure 1:**
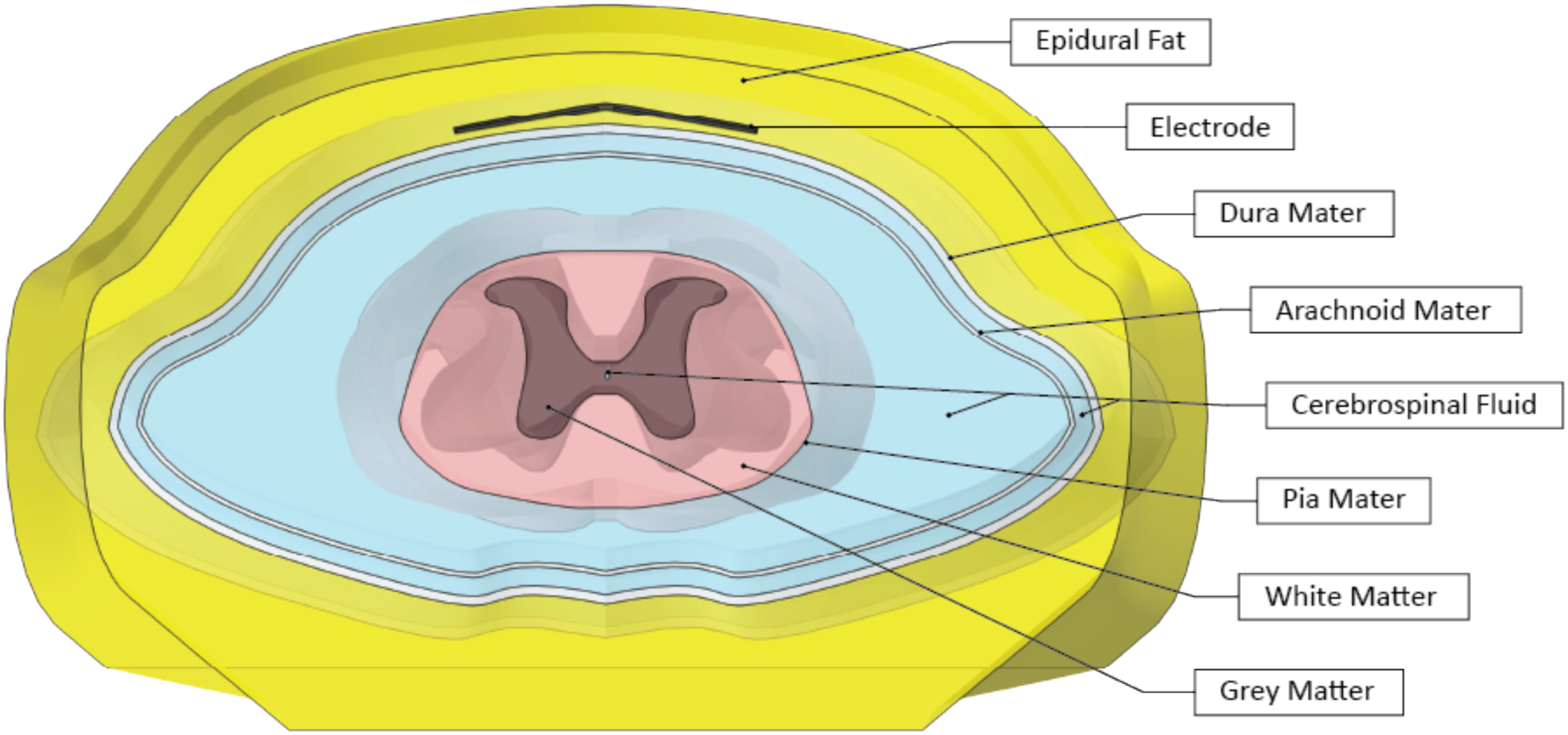
Computational Model Domains. The biophysically accurate layers of the rodent spinal cord were comprised of materials representing epidural fat, platinum electrode faces, dura mater, arachnoid mater, cerebrospinal fluid, pia mater, white matter, and grey matter.

**Supplementary Figure 2:**
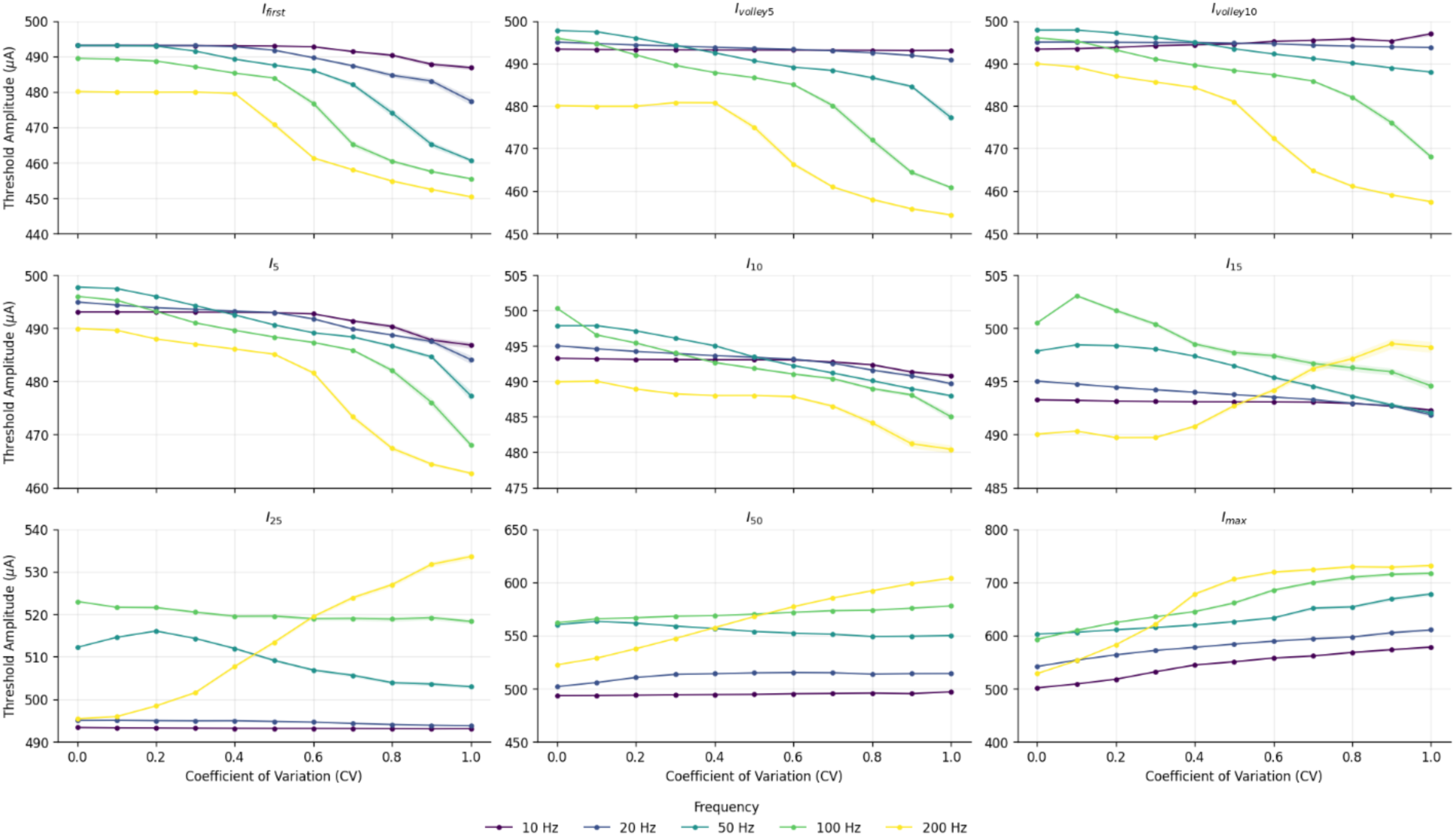
Spike-Count Threshold Amplitudes. Plots show the simulated input current (µA) required to satisfy the different spike-count criteria as a function of CV. Colors correspond to frequencies of 10, 20, 50, 100, and 200 Hz as described by the bottom legend. Datapoints at each CV value represent the mean ± SEM across 100 simulated repetitions.

**Supplementary Figure 3:**
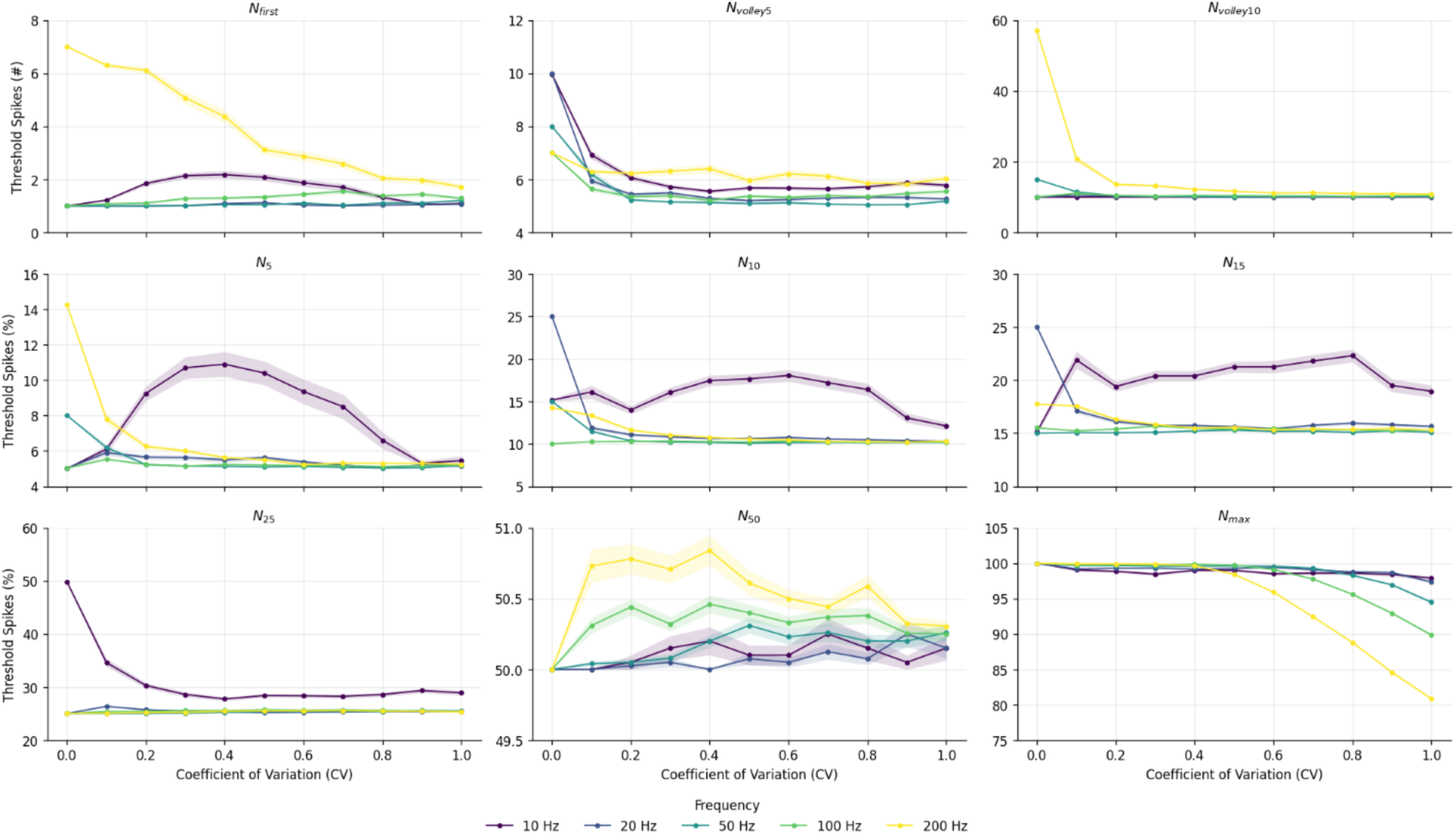
Regime Spike Counts. Plots show the number (top row) or percentage (bottom two rows) of simulated criteria-satisfying spikes produced as a function of CV. Colors correspond to frequencies of 10, 20, 50, 100, and 200 Hz as described by the bottom legend. Datapoints at each CV value represent the mean ± SEM across 100 simulated repetitions.

**Supplementary Figure 4:**
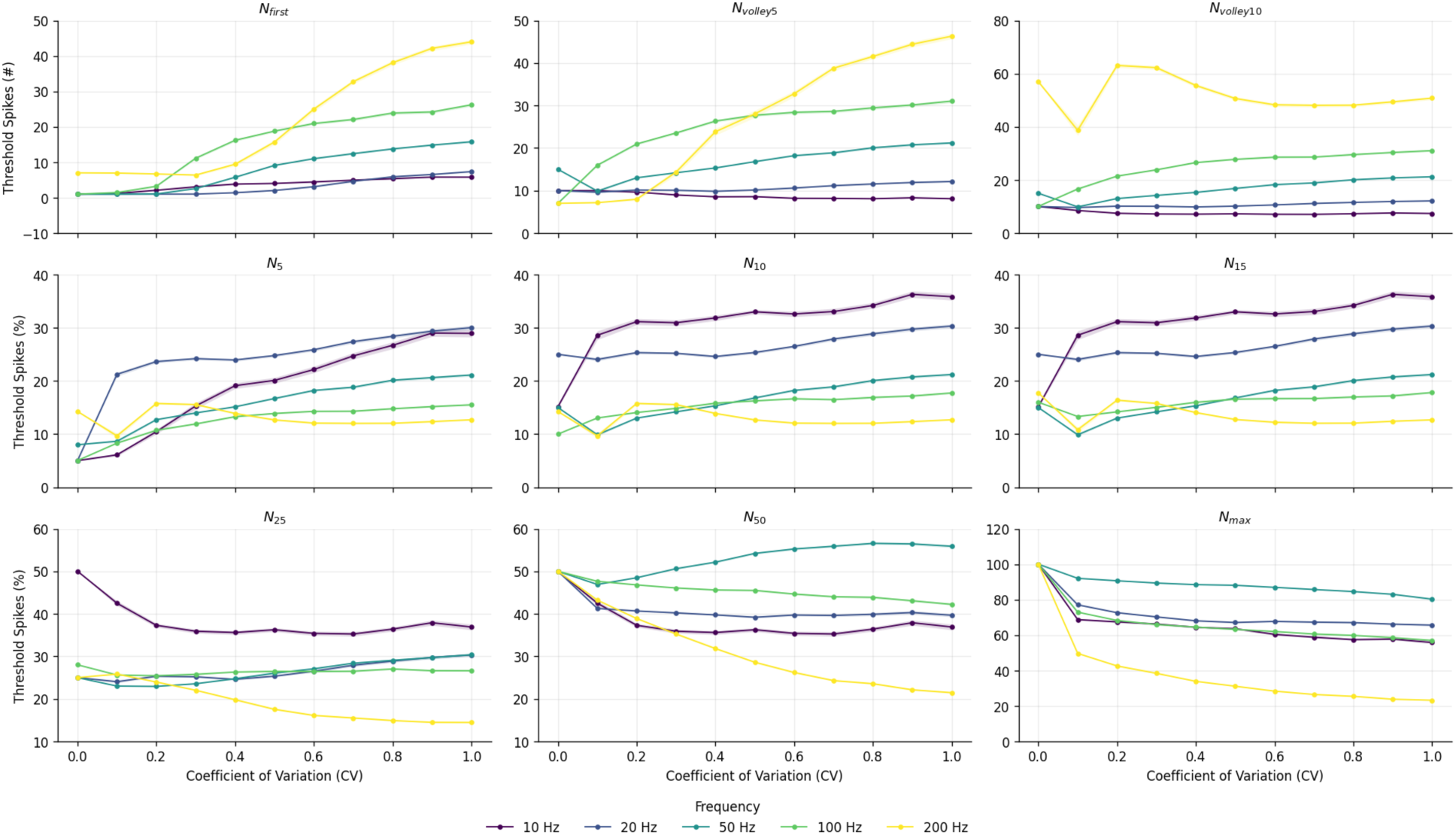
CV0-Based Regime Spike Counts. Plots show the number (top row) or percentage (bottom two rows) of simulated spikes produced as a function of CV using the spike-count threshold amplitudes determined at CV=0 for all CV values. Colors correspond to frequencies of 10, 20, 50, 100, and 200 Hz as described by the bottom legend. Datapoints at each CV value represent the mean ± SEM across 100 simulated repetitions.

**Supplementary Figure 5:**
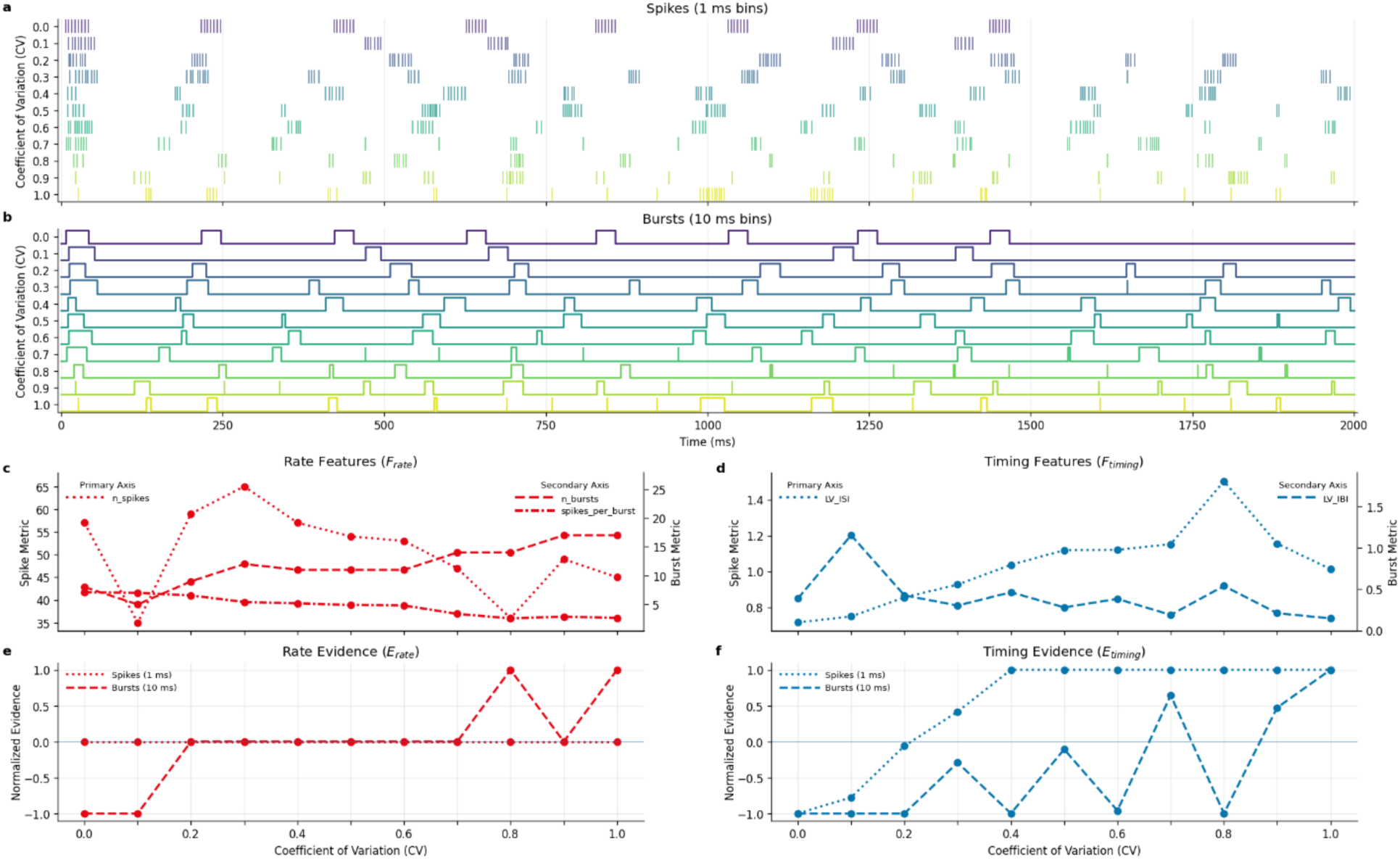
Feature Engineering. This figure shows an illustrative example of the process used to convert spike trains into signed template evidence relative to the CV = 0 and CV = 1 endpoint templates for a single 200 Hz repetition and the CV0-based *I*_10_ spike-count threshold amplitude. **a)** Spikes simulated for all CVs using the CV0-based *I*_10_ input current amplitude. **b)** Bursts for all CVs computed by collapsing contiguous active 10-ms bins into single events. **c)** Rate features (*F*_*rate*_) for spikes ( *n*_*spikkes*_) and bursts (*n*_*bursts*_, spik*kes*_*per*_*burst*) plotted on the primary and secondary y-axes, respectively, as functions of CV. **d)** Timing features (*F*_*timing*_) for spikes (*LV*_*IBI*_) and bursts (*LV*_*IBI*_) plotted on the primary and secondary y-axes, respectively, as functions of CV. **e)** Rate evidence (*E*_*rate*_) for spikes and bursts computed for each CV by comparing feature differences between CV = 0 and CV = 1 endpoint templates and satisfying the JND ratio > 2.5. **f)** Timing evidence (*E*_*timing*_) for spikes and bursts computed for each CV by comparing feature differences between CV = 0 and CV = 1 endpoint templates.

**Supplementary Figure 6.**
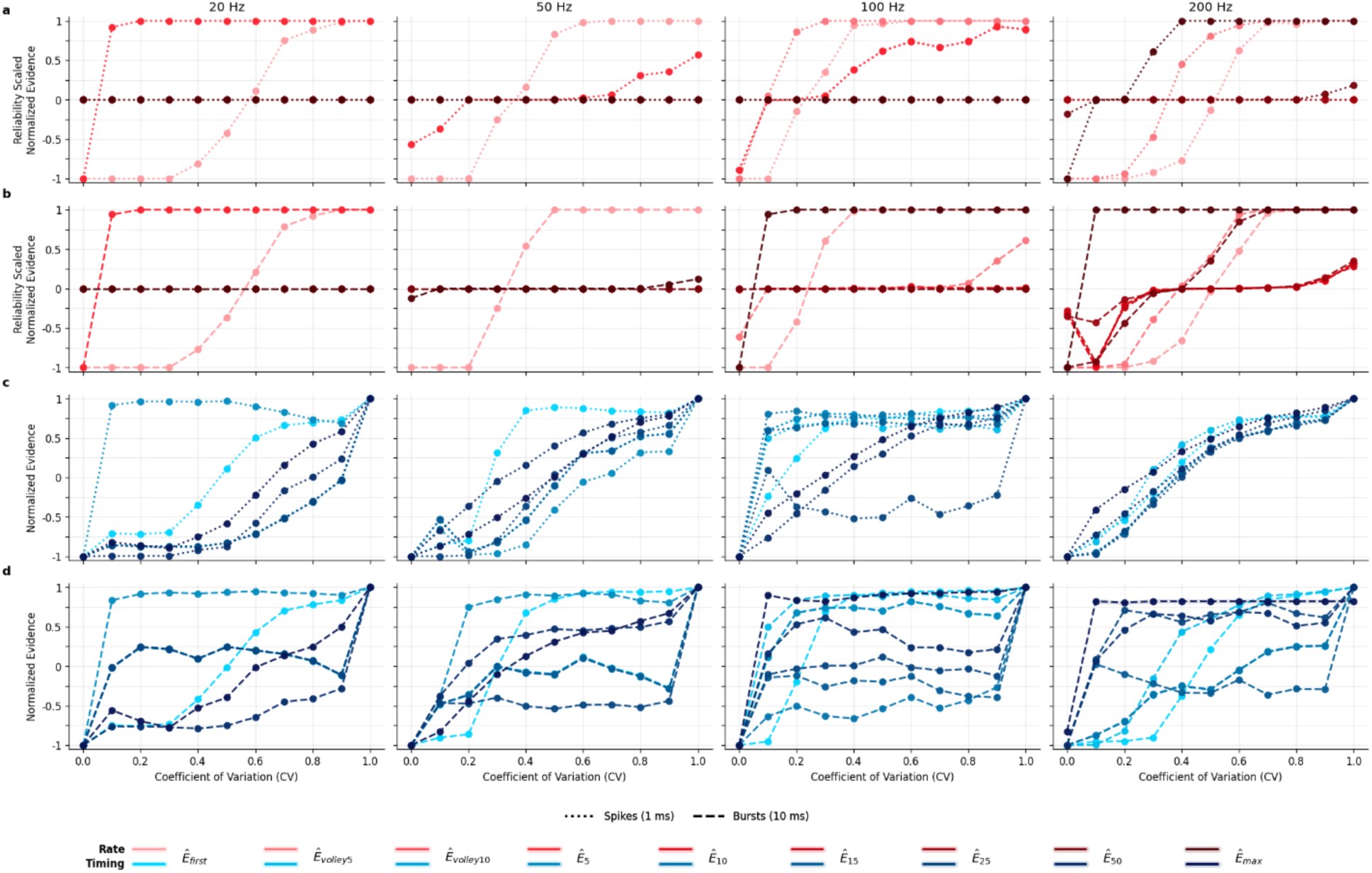
Normalized Evidence Curves. This figure shows the signed rate (*Ê*_*rate*_) and timing (*Ê*_*timing*_) template evidence for the spike and burst features for each CV0-based spike-count regime. Datapoints are mean ± SEM across the 200 Monte Carlo training sets, and the negative and positive evidence values correspond to CV = 0 and CV = 1, respectively. **a)** Reliability scaled spike rate evidence as a function of CV. **b)** Reliability scaled burst rate evidence as a function of CV. **c)** Spike timing evidence as a function of CV. **d)** Burst timing evidence as a function of CV.

**Supplementary Figure 7:**
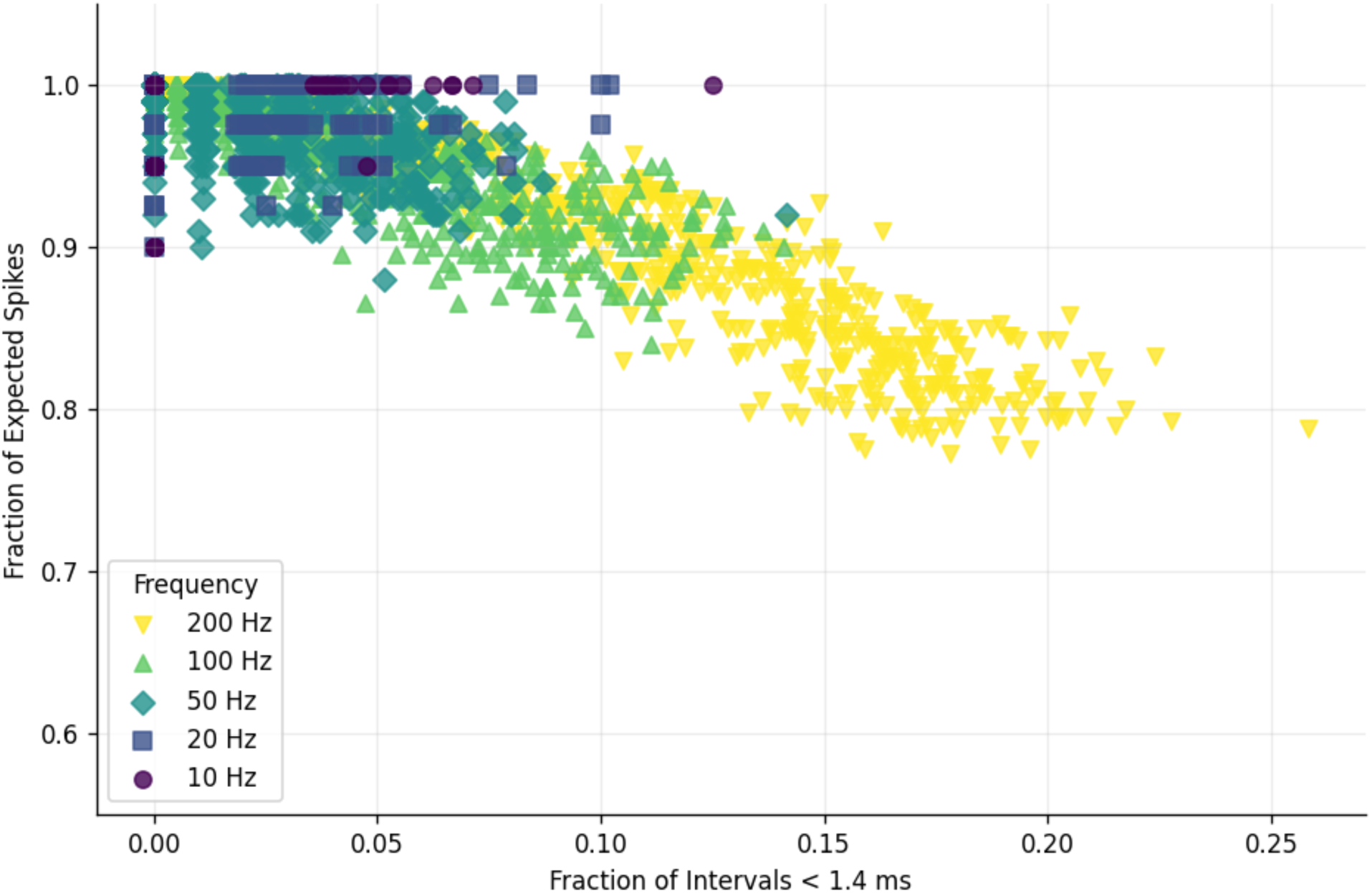
Sub-Refractory Inter-Pulse Intervals. This figure shows the fraction of expected spikes (*N*_*max*_⁄(2 ∗ *freq*)) at the plateau input current (*I*_*max*_) as a function of the fraction of inter-pulse intervals < 1.4 ms. Colors and markers described in the bottom legend represent frequencies 10, 20, 50, 100, and 200 Hz. Each datapoint represents a single simulated repetition.

**Supplementary Figure 8:**
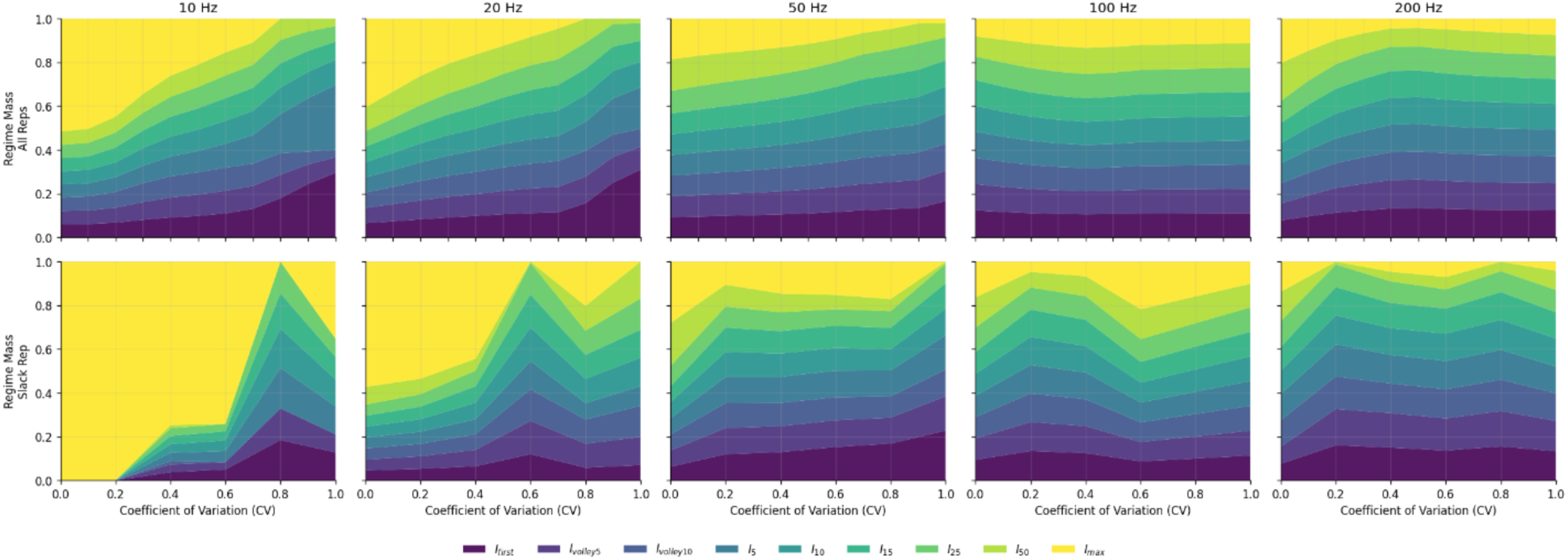
Detection Threshold Spike-Count Regime Mixtures. This figure shows stacked area plots for the per-point mixtures across frequencies and CVs identified for the behavioral quadratic fit using all repetitions (top row) and the behavioral datapoint fit using the Slack-matched repetition (bottom row). Colors represent the different spike-count thresholds, as described by the bottom legend.

**Supplementary Figure 9:**
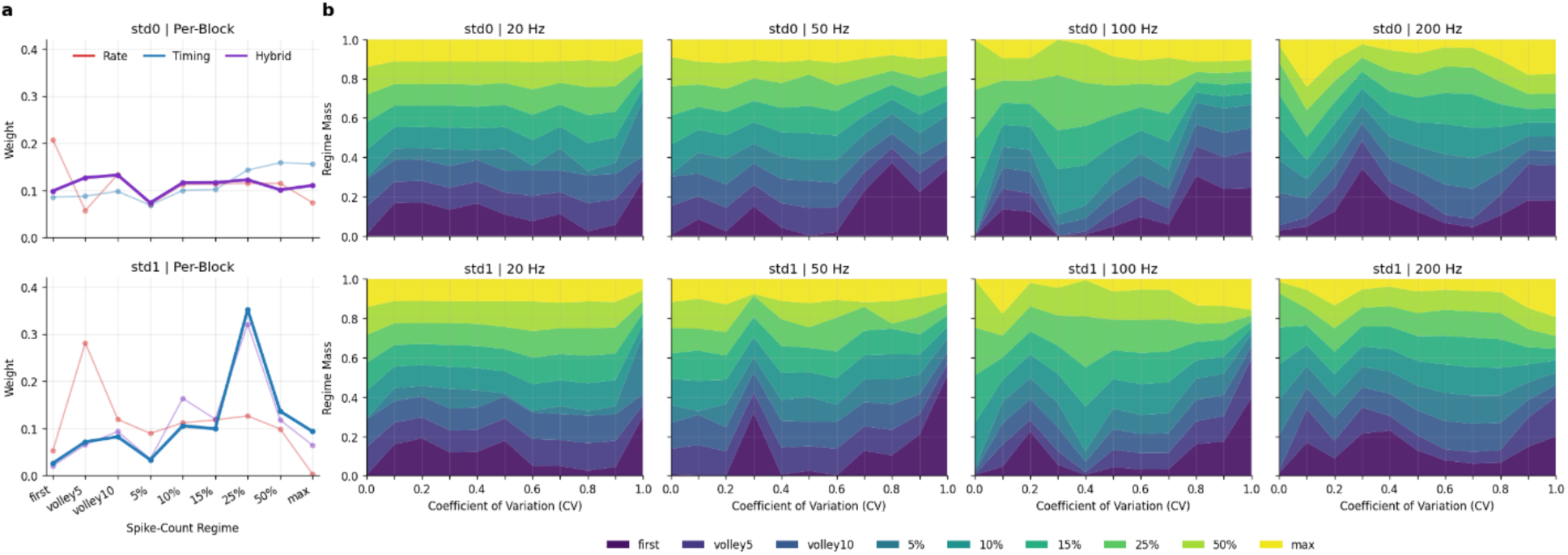
Periodicity Discrimination Spike-Count Regime Mixtures. This figure shows the learned CV0-based spike-count regime mixture weights for the per-block and per-point models. The top and bottom rows show std0 and std1 blocks, respectively. **a)** Block-shared regime weights for the rate (red), timing (blue), and hybrid (purple) per-block models; the best-fit model is shown in bold. **b)** Stacked area plots for the per-point mixtures across frequencies and CVs; colors represent the different CV0-based spike-count regimes, as described by the bottom legend.

**Supplementary Table 1:** Spinal Cord Material Properties. This table shows the conductivities (S/m) used for the COMSOL spinal cord anatomical structures and stimulation electrode.

| Material Name | Conductivity (S/m) |
| --- | --- |
| Epidural Fat | 4e-2 |
| Electrode | 9.4e6 (Pt faces), 1e-12 (PDMS insulator) |
| Dura Mater | 5.1e-2 |
| Arachnoid Mater | 5.1e-2 |
| Cerebrospinal Fluid | 1.7 |
| White Matter | 6e-1 (Longitudinal), 8.3e-2 (Transverse) |
| Grey Matter | 2.3e-1 |
| Pia Mater | 5.1e-2 |

**Supplementary Table 2:** Discrimination Behavioral Matching Performance. This table shows the std0 and std1 block-averaged test RMSE values for the rate, timing, and hybrid observer models using per-block mixtures. Block test RMSE values are shown as mean ± SD, and the mean difference vs. best column are shown as percentages. Asterisks indicate statistical significance from paired Wilcoxon tests across Monte Carlo draws (*** p < 0.001).

| Block | Model | Block Test RMSE (Mean ± SD) | Mean Difference vs. Best (%) |
| --- | --- | --- | --- |
| std0 | hybrid | 0.074 ± 0.004 | 0 |
| std0 | timing | 0.106 ± 0.004*** | 42.36 |
| std0 | rate | 0.108 ± 0.002*** | 45.68 |
| std1 | timing | 0.094 ± 0.007 | 0 |
| std1 | hybrid | 0.097 ± 0.008*** | 3.03 |
| std1 | rate | 0.154 ± 0.005*** | 63.82 |

